# A newly designed GABA-AT inactivator, (*S*)-MeCPP-115, suppresses paclitaxel-induced neuropathic pain in mice

**DOI:** 10.1101/2025.10.20.683456

**Authors:** Luana Assis Ferreira, Koon Mook Kang, Ifeoluwa Solomon, Richard B. Silverman, Andrea G. Hohmann

**Affiliations:** Department of Psychological and Brain Sciences, Indiana University, Bloomington, IN 47405; Gill Institute for Neuroscience, Indiana University, Bloomington, IN 47405; Program in Neuroscience, Indiana University, Bloomington, IN 47405; Department of Chemistry, Northwestern University, Evanston, IL 60208; Department of Molecular Biosciences, Northwestern University, Evanston, IL 60208; Chemistry of Life Processes Institute, Northwestern University, Evanston, IL

**Keywords:** Chemotherapy-induced peripheral neuropathy, paclitaxel, GABA-aminotransferase inhibition, (S)-MeCPP-115, neuropathic pain

## Abstract

**Background:** Chemotherapy-induced peripheral neuropathy (CIPN) is a common and debilitating side effect of paclitaxel treatment. Pharmacological strategies that enhance inhibitory tone, via inhibition of γ-aminobutyric acid aminotransferase (GABA-AT), the enzyme which degrades endogenous GABA, provides analgesic benefit in preclinical studies. (*S*)-MeCPP-115 is a novel selective GABA-AT inactivator designed from CPP-115 to minimize off-target activity. Whether (*S*)-MeCPP-115 shows efficacy in preclinical pain models is unknown.

**Methods:** *In vitro* assays of cell viability were conducted to ascertain whether (*S*)-MeCPP-115 interfered with antitumor activity of paclitaxel or produced cytotoxicity in normal cells. Male mice received paclitaxel to induce chemotherapy-induced peripheral neuropathy *in vivo*. In mice with established paclitaxel-induced mechanical hypersensitivity, (*S*)-MeCPP-115 was administered via acute and chronic administration using intraperitoneal (i.p.) or intrathecal (i.t.) dosing strategies. Mechanical sensitivity was assessed in all mice with an electronic von Frey analgesiometer before and after paclitaxel and pharmacological treatments. Locomotor activity was also measured in the same subjects to assess possible motor impairment.

**Results:** In MTT assays, (*S*)-MeCPP-115 did not alter the cytotoxic activity of paclitaxel in 4T1 breast cancer cells and did not produce cytotoxicity in non-tumor HEK293 cells. (*S*)-MeCPP-115 reduced paclitaxel-induced mechanical hypersensitivity after i.p. and i.t. administration. Therapeutic efficacy was maintained with repeated dosing without development of tolerance. (*S*)-MeCPP-115 did not alter paw withdrawal thresholds in mice that received the Cremophor-based vehicle in lieu of paclitaxel following acute or chronic dosing. Systemic treatment with (*S*)-MeCPP-115 was well tolerated, whereas i.t. administration produced only minimal locomotor effects.

**Conclusion:** (*S*)-MeCPP-115 did not interfere with the ability of paclitaxel to produce tumor cell cytotoxicity *in vitro* and did not produce cytotoxicity in non-tumor cells. Systemic and intrathecal administration of (*S*)-MeCPP-115 produced robust suppression of mechanical hypersensitivity in a mouse model of paclitaxel-induced CIPN without major adverse effects. Selective GABA-AT inhibition represents a promising therapeutic approach for suppressing paclitaxel-induced mechanical hypersensitivity. Further preclinical characterization of the therapeutic profile of (*S*)-MeCPP-115 is warranted.

## Introduction

Chemotherapy-induced peripheral neuropathy (CIPN) is a frequent and often debilitating complication of cancer therapy, occurring in up to 90% of patients receiving taxane-based treatment such as paclitaxel [1]. Characteristic symptoms include painful sensory disturbances and mechanical hypersensitivity. Mounting evidence implicates dysregulation of inhibitory neurotransmission in CIPN. Signaling deficits in the inhibitory neurotransmitter gamma-aminobutyric acid (GABA) within central and spinal nociceptive circuits contribute to the development and maintenance of neuropathic pain [2]. CIPN has been associated with reduced GABAergic inhibition in the anterior cingulate cortex, thalamus, insula, and spinal cord [3, 4]. Consistently, pharmacological or cellular strategies aimed at restoring spinal GABAergic tone, such as GABA transporter blockade or transplantation of GABAergic precursors attenuate paclitaxel-evoked hypersensitivity in rodent models. These findings support the concept that augmenting GABAergic neurotransmission may provide analgesic benefit in CIPN [5, 6]. One indirect approach to enhancing GABAergic signaling is inhibition of γ-aminobutyric acid aminotransferase (GABA-AT), the enzyme responsible for GABA degradation. The first-generation GABA-AT inhibitor vigabatrin demonstrated efficacy in epilepsy and substance use disorders, but its clinical use was severely limited by off-target toxicities, including irreversible visual field defects [1, 7, 8]. To overcome these limitations, second-generation inhibitors were developed with improved pharmacological properties. Among them, CPP-115 was rationally designed to increase efficiency and reduce off-target activity compared to vigabatrin. Preclinical studies demonstrated robust anticonvulsant efficacy, and CPP-115 successfully completed a Phase I clinical trial, with subsequent compassionate use in patients with infantile spasms [9–11]. OV329, a next-generation GABA-AT inhibitor, has completed Phase 1 evaluation in healthy volunteers, demonstrating robust target engagement, favorable safety and tolerability, and 100-to 1,000-fold greater potency than vigabatrin in preclinical studies [12, 13]. Our group recently showed that OV329 produced antinociceptive effects in both CFA-and paclitaxel-induced pain models, was not self-administered, and did not induce conditioned place preference [6]. Building on these findings, (*S*)-MeCPP-115 was developed as a novel GABA-AT inactivator with enhanced selectivity and reduced off-target activity at ornithine aminotransferase (OAT), aiming to minimize off target effects while preserving analgesic efficacy. In the present study, we first evaluated (*S*)-MeCPP-115 with paclitaxel in cell-based assays to confirm that (*S*)-MeCPP-115 does not interfere with the antitumor activity of paclitaxel and does not produce cytotoxicity on its own. Based on these findings, we assessed the efficacy of (*S*)-MeCPP-115 in a mouse model of paclitaxel-induced neuropathic pain. Intraperitoneal (i.p.) and intrathecal (i.t.) routes of administration were tested to determine the ability of (*S*)-MeCPP-115 to reverse established mechanical hypersensitivity. In addition, locomotor activity was evaluated to rule out potential motor impairment, which is relevant to the interpretation of antiallodynic efficacy.

## Materials and Methods

### Cell Viability Assay

Methods are similar to those used in our previously published work [6]. Briefly, 4T1 murine breast cancer cells were maintained in RPMI-1640 medium, and human embryonic kidney HEK293 cells in DMEM, both supplemented with 10% fetal bovine serum and 1% penicillin–streptomycin. Cells were cultured at 37 °C in a humidified incubator with 5% CO₂. Cell viability was determined using the MTT assay (Roche, Indianapolis, IN) according to the manufacturer’s protocol, as previously described. Half-maximal inhibitory concentrations (IC₅₀) were calculated by nonlinear regression analysis. Drug–drug interaction analyses were performed using Combenefit (Cancer Research UK Cambridge Institute, Cambridge, UK) and SynergyFinder (https://synergyfinder.fimm.fi).

Three reference models were applied: (i) Highest Single Agent (HSA), which assumes the expected combination effect equals the maximal effect of either drug alone at the same concentrations; (ii) Bliss Independence, which assumes independent drug action with the expected effect derived from the probabilistic combination of single-agent responses; and (iii) Loewe Additivity, which assumes both drugs act through a shared mechanism and estimates the expected effect accordingly.

### Animals

All experimental procedures were approved by the Institutional Animal Care and Use Committee of Indiana University Bloomington and conducted in accordance with the National Institutes of Health Guide for the Care and Use of Laboratory Animals. Male CD1 mice (7-8 weeks old; Jackson Laboratories, Bar Harbor, ME) were housed in groups under a standard 12 h light/dark cycle with *ad libitum* access to food and water. In all experiments, investigators were blinded to treatment condition.

### Drug Preparation and Administration

Paclitaxel (Tecoland Corporation, Irvine, CA) was prepared in a vehicle composed of Cremophor EL (Sigma-Aldrich, St. Louis, MO), ethanol, and 0.9% saline in a 1:1:18 (v/v/v) ratio. (*S*)-MeCPP-115 was synthesized by KMK in the laboratory of Richard B. Silverman (Department of Chemistry, Northwestern University, USA) and dissolved in sterile 0.9% saline. For intrathecal (i.t.) administration, (*S*)-MeCPP-115 was diluted in 0.9% saline containing 6% glucose and delivered in a total volume of 5 µL. Injections were performed without anesthesia at the L5–L6 intervertebral space, using the tail-flick reflex as confirmation of correct intrathecal placement. Intraperitoneal (i.p.) injections were administered at a volume of 10 mL/kg.

### Assessment of Mechanical Sensitivity

Mice were acclimated for 1h on an elevated metal mesh platform prior to testing. Mechanical paw withdrawal thresholds were measured using a von Frey anesthesiometer (Almemo 2450 with 90 g probe) applied to the plantar surface of the hind paw. Each paw was tested in duplicate, and the average value from each animal was used for analysis.

### Paclitaxel-Induced Peripheral Neuropathy

Peripheral neuropathy was induced by four i.p. injections of paclitaxel (4 mg/kg per injection; cumulative dose 16 mg/kg) or Cremophor-based vehicle on days 0, 2, 4, and 6, as previously described. (*S*)-MeCPP-115 (10 mg/kg, i.p. or 100 µg, i.t.) or saline was administered after neuropathy was established and stable, 15 days following the first paclitaxel injection. Mechanical thresholds were measured at baseline and at the indicated time-points after (*S*)-MeCPP-115 or saline administration. In chronic systemic dosing studies (*S*)-MeCPP-115 (10 mg/kg, i.p. per day) or saline was administered once daily for 9 consecutive days in mice that were treated previously with paclitaxel or its vehicle. In chronic intrathecal dosing studies, (*S*)-MeCPP-115 (100 µg per day, i.t.) or vehicle was administered once daily for 7 consecutive days in mice that previously received either paclitaxel or its cremophor-based vehicle.

### Locomotor Activity

Locomotor behavior was evaluated using automated activity meters (Omnitech, Columbus, OH). Mice were initially acclimated to the testing room before assessment, and activity was recorded for 30 min beginning 1 h after (*S*)-MeCPP-115 administration. Parameters analyzed included: total distance traveled (overall locomotor output), horizontal activity (beam breaks reflecting exploratory movement), rest time (periods of inactivity ≥1 s), movement time (cumulative time engaged in locomotion), time spent in the center of the chamber (index of anxiety-like behavior), and ambulatory velocity (average speed while active). Assessments were conducted on day 9 for the i.p. cohort and on day 7 for the i.t. cohort.

## Statistical Analysis

Cell viability data were analyzed with Combenefit and SynergyFinder according to the Bliss Independence, HSA, and Loewe Additivity reference models, as previously described [6]. Behavioral data were analyzed by One-way or Two-way ANOVA. Post hoc comparisons were subsequently performed using Sidak’s test for two-group comparisons or Tukey’s test for multiple comparisons among four groups, using GraphPad Prism 10 (GraphPad Software, La Jolla, CA). A value of p < 0.05 was considered statistically significant.

## Results

### Paclitaxel-induced cytotoxicity in 4T‘1 cells is preserved in the presence of (*S*)-MeCPP-115

Paclitaxel significantly reduced the viability of 4T1 cells, IC₅₀ of 15.18 nM (95% CI: 66.90–91.48 nM) (**Fig. 1A**). In contrast, (*S*)-MeCPP-115 alone had no effect on 4T1 viability (undetermined IC₅₀ (95% CI): 102.3–113.9 µM) (**Fig. 1B**). In HEK293 cells, paclitaxel modestly decreased cell viability at higher concentrations, with an IC₅₀ ranging from 83.73 to 103.2 nM (**Fig. 1C**). (*S*)-MeCPP-115 showed no measurable effect, with an undetermined IC₅₀ (95% CI: 93.99–107.1 µM) (**Fig. 1D**). These data represent means with 95% confidence intervals from three independent MTT experiments for each cell line. Computational quantification of drug combination effects across a broad range of molar ratios revealed that paclitaxel in combination with (*S*)-MeCPP-115 did not induce antagonism in 4T1 cells under the Bliss independence model (**Fig. 2E**), Highest Single Agent (HSA) model (**Fig. 2F**), or Loewe additivity model (**Fig. 2G**). In HEK293 cells, paclitaxel modestly impacted cell viability (**Fig. 3A**) whereas (*S*)-MeCPP-115 (**Fig. 3B**) failed to do so and neither compound altered the dose–response curve of the other (**Fig. 3D**).

**Figure 1.**
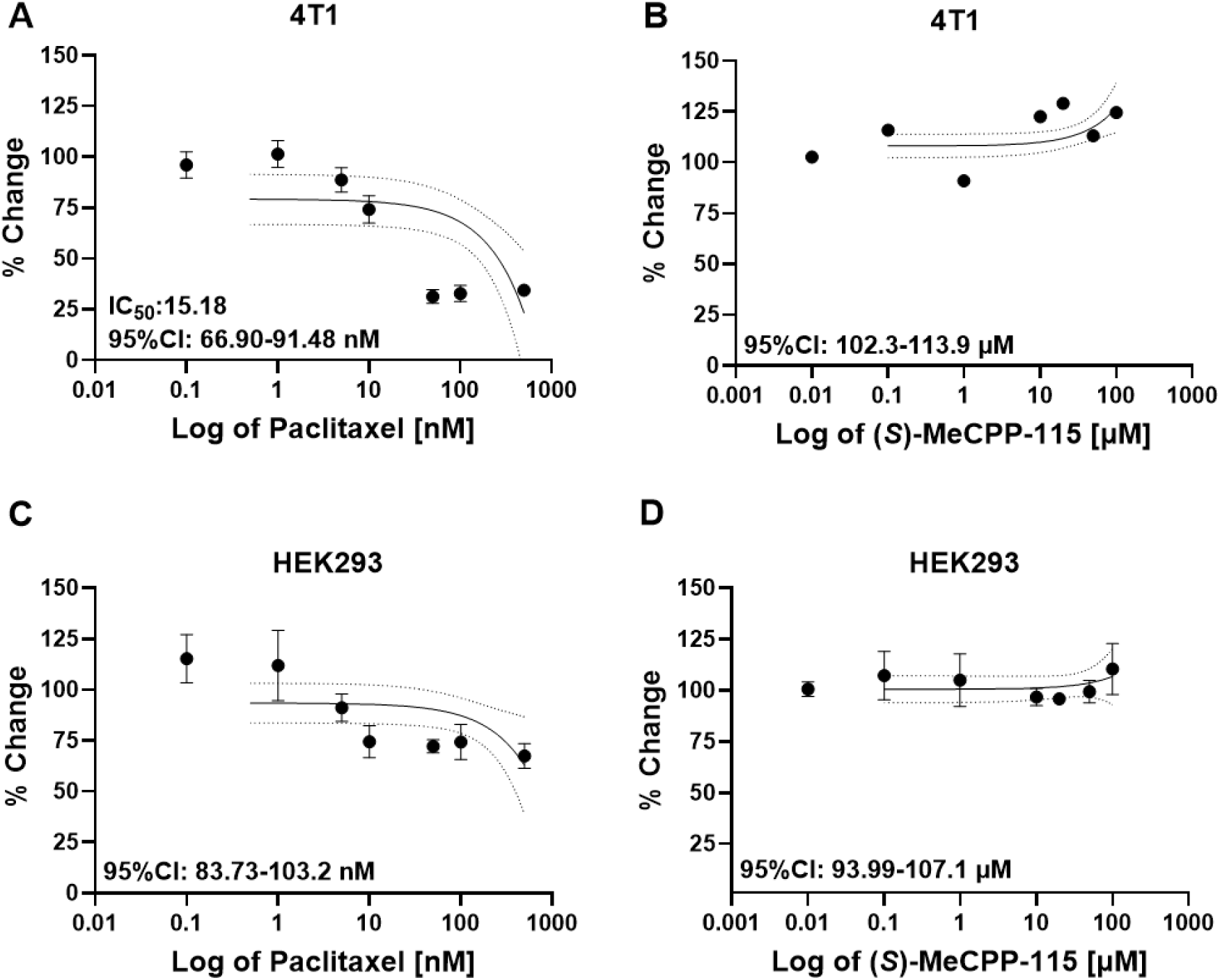
Impact of paclitaxel and (*S*)-MeCPP-115 on cell viability in 4T1 and HEK293 cells. Dose–response curves of 4T1 cells treated with increasing concentrations of paclitaxel (**A**) or (*S*)-MeCPP-115 (**B**). Paclitaxel showed a potent cytotoxic effect with an IC₅₀ of 15.18 nM (95% CI: 6.69–91.48 nM), indicating high sensitivity of tumor cells to the treatment. In contrast, (*S*)-MeCPP-115 did not display significant cytotoxicity, with an IC₅₀ above 100 µM (95% CI: 102.3–113.9 µM). (**C**, **D**) Corresponding curves for HEK293 cells showed no appreciable cytotoxic effects for either paclitaxel (IC₅₀: 83.73–103.2 nM) or (*S*)-MeCPP-115 (IC₅₀: 93.99–107.1 µM), suggesting selectivity of paclitaxel toward tumor cells. Data are expressed as mean ± SEM from at least three independent experiments.

**Figure 2.**
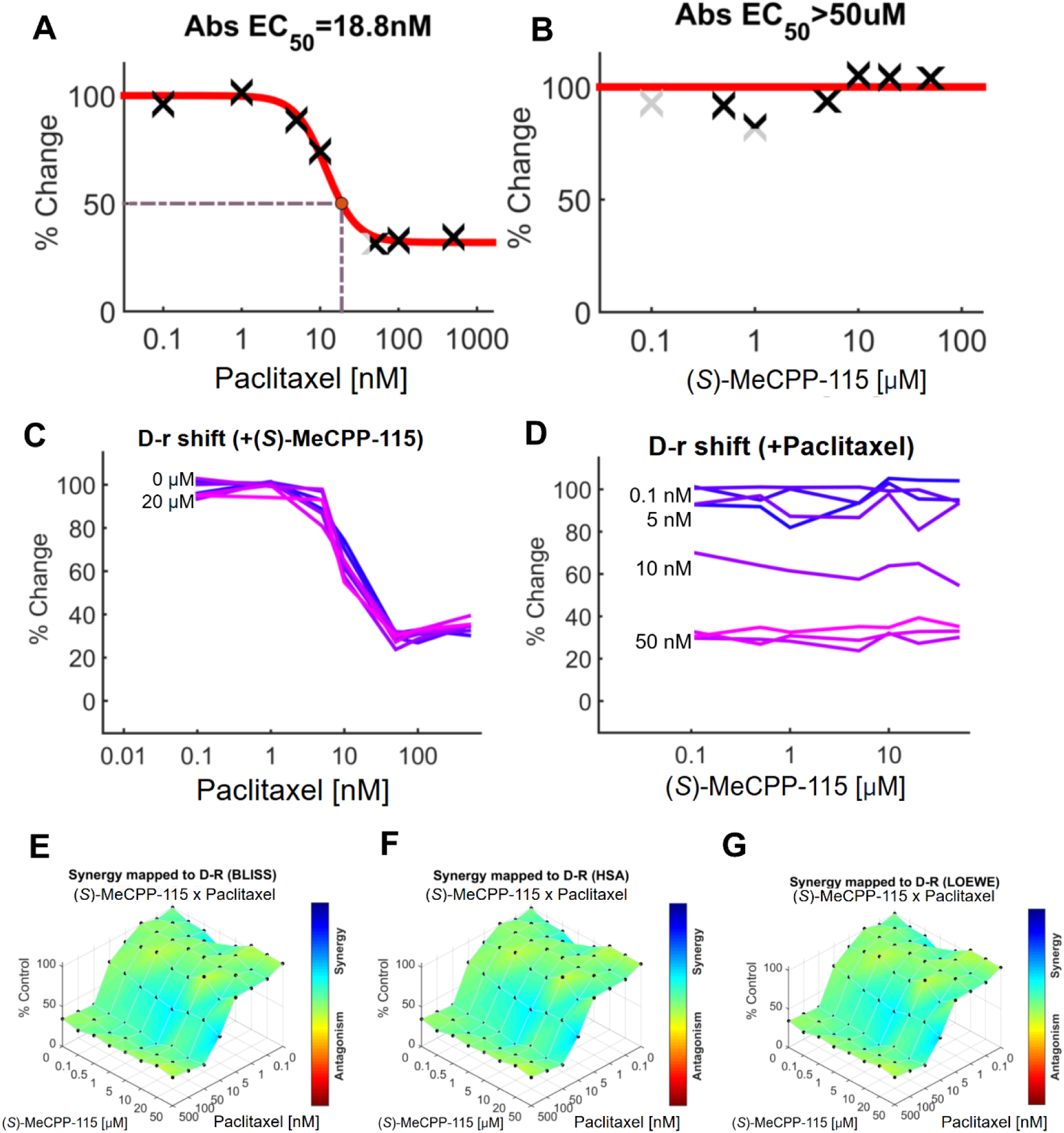
(*S*)-MeCPP-115 does not interfere with the cytotoxicity of paclitaxel in reducing breast cancer cell viability in murine 4T1 cells. Paclitaxel alone reduced 4T1 cell viability with an EC₅₀ of 18.8 nM (**A**). (*S*)-MeCPP-115 alone had no significant effect on tumor cell viability (EC₅₀ > 50 µM) (**B**). Single-agent and combination responses indicate that the paclitaxel dose–response curve slightly shifted in the presence of increasing concentrations of (*S*)-MeCPP-115 (**C**). Similarly, the (*S*)-MeCPP-115 dose–response curve was not affected by the presence of paclitaxel (**D**). Three-dimensional plots display the full matrix of dose–response interactions, with blue regions indicating synergy and red regions indicating antagonism. Three synergy models — Bliss (**E**), Highest Single Agent (HSA, **F**), and Loewe (**G**) — highlight small areas of synergy between paclitaxel and (*S*)-MeCPP-115 in this cell line. Cell viability is expressed as percent of control (wells incubated only with corresponding medium). Each experiment was performed in triplicate (n = 3). Data shown represent the composite fit derived from all three MTT assay plates.

**Figure 3.**
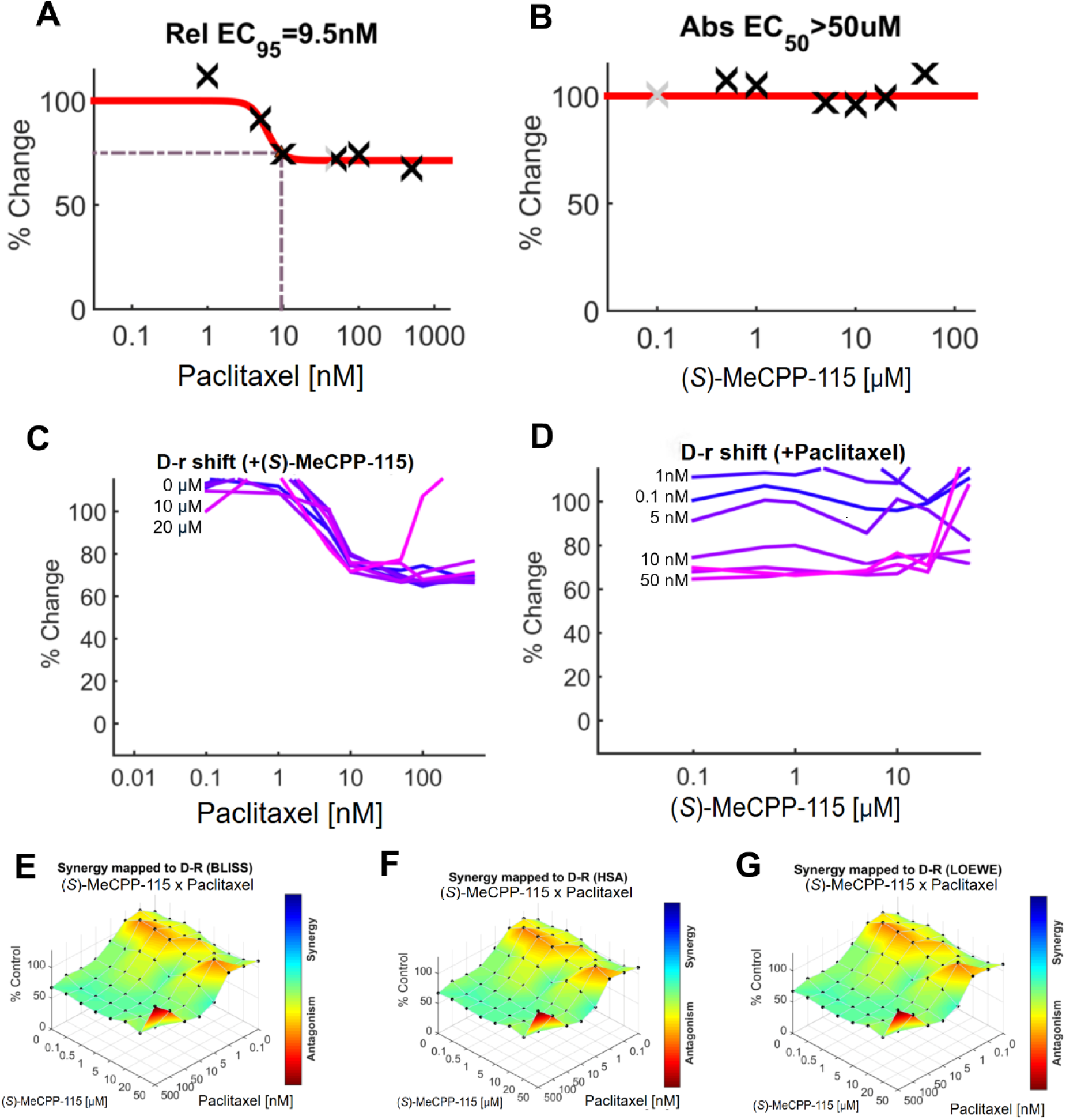
(*S*)-MeCPP-115 does not induce cytotoxicity in non-cancerous HEK293 cells. Paclitaxel had little effect on HEK293 cell viability, with a relative EC₉₅ of 9.5 nM (**A**), whereas (*S*)-MeCPP-115 alone had no effect on cell viability (EC₅₀ > 50 µM) (**B**). Single-agent and combination responses indicate that neither agent alone nor in combination markedly affected the viability of HEK293 cells (**C**, **D**). Three-dimensional plots display the full matrix of dose–response interactions, with blue regions indicating synergy and red regions indicating antagonism. Three synergy models — Bliss (**E**), Highest Single Agent (HSA, **F**), and Loewe (**G**) — all suggest little to no effect of paclitaxel, (*S*)-MeCPP-115, or their combination on HEK293 cell viability. Cell viability is expressed as percent of control (wells incubated only with corresponding medium). Each experiment was performed in triplicate (n = 3). Data shown represent the composite fit derived from all three MTT assay plates.

Computational quantification of drug combination effects across a broad range of molar ratios revealed that paclitaxel in combination with (*S*)-MeCPP-115 did not induce antagonism in 4T1 cells under the Bliss independence model (**Fig. 3E**), Highest Single Agent (HSA) model (**Fig. 3F**), or Loewe additivity model (**Fig. 3G**).

Consistent with these observations, quantitative synergy analysis using the SynergyFinder platform (**Table 1**) indicated that the combination of paclitaxel with (*S*)-MeCPP-115 yielded values within the range of weak or negligible synergy in 4T1 cells, with no evidence of antagonism. In HEK293 cells, synergy scores further confirmed the absence of any interaction between the two agents.

**Table 1.** Synergy scores for the combination of paclitaxel and (*S*)-MeCPP-115 on cell viability.

| Cell Line | Method | Synergy Score | Most Synergistic Area Score |
| --- | --- | --- | --- |
| 4T1 | BLISS | -4.893 ± 4.64 | 3.939 |
|  | LOEWE | -7.966 ± 4.64 | 6.907 |
|  | HSA | 0.634 ± 4.64 | 7.508 |
| HEK293 | BLISS | -10.463 ± 5.96 | -3.633 |
|  | LOEWE | -11.917 ± 5.96 | -1.429 |
|  | HSA | -3.274 ± 5.96 | 1.334 |

Scores greater than 10 are considered synergistic. Scores between -10 and 10 are likely additive. Scores less than -10 suggest evidence for antagonism.

### Repeated i.p. administration of (*S*)-MeCPP-115 reduces paclitaxel-evoked mechanical hypersensitivity without inducing motor side effects

The experimental timeline illustrated in **Figure 4A** shows induction of neuropathic pain by paclitaxel (PAX) treatment, followed by 9 daily injections of (*S*)-MeCPP-115 or vehicle (i.p.) and timing of the behavioral assessments throughout the course of treatment. Paclitaxel treatment produced robust and persistent mechanical hypersensitivity compared with vehicle controls (**Fig. 4B**; treatment: F(4,110)=28.78, p<0.0001; time: F(4,110)=37.46, p<0.0001; interaction: F(1,110)=240.4, p<0.0001), with significant decreases in paw withdrawal threshold observed from day 7 to day 15 (p<0.0001) following initiation of paclitaxel dosing. Acute administration of (*S*)-MeCPP-115 (10 mg/kg, i.p.) reduced mechanical hypersensitivity for over 2 h, an effect that dissipated by 24 h (**Fig. 4C**; treatment: F(1,60)=43.19, p<0.0001; time: F(5,60)=7.213, p<0.0001; interaction: F(5,60)=2.441, p=0.0443). (*S*)-MeCPP-115 (i.p.) suppressed paclitaxel-induced mechanical hypersensitivity at 0.5, 1, and 2 h post-injection (p<0.05). In vehicle-treated controls, (*S*)-MeCPP-115 did not alter mechanical thresholds during the acute phase (**Fig. 4D**; treatment: F(1,60)=3.471, p=0.0673; time: F(5,60)=1.004, p=0.4233; interaction: F(5,60)=0.3620, p=0.8725). Repeated once daily (i.p.) administration of (*S*)-MeCPP-115 (20 mg/kg, i.p. per day) reversed paclitaxel-evoked hypersensitivity when assessed 1h post-injection, with no evidence of tolerance observed across the 9-day treatment period (**Fig. 4E**; treatment: F(1,100)=178.3, p<0.0001; time: F(9,100)=2.846, p=0.0050; interaction: F(9,100)=1.223, p=0.2896). In control mice that received vehicle in lieu of paclitaxel, repeated (10 mg/kg, i.p. per day) dosing had no effect on mechanical thresholds, which remained near baseline (**Fig. 4F**; treatment: F(1,100)=1.447, p=0.2318; time: F(9,100)=1.963, p=0.0515; interaction: F(9,100)=2.210, p=0.0273). At 24 h post-injection, (*S*)-MeCPP-115 did not produce a residual antinociceptive effect (**Fig. 4G**; treatment: F(1,90)=2.240, p=0.1380; time: F(8,90)=1.397, p=0.2085; interaction: F(8,90)=1.942, p=0.0632). Similarly, in controls mice receiving the Cremophor based vehicle, paw withdrawal thresholds remained unchanged over time with the except for a transient decrease on day 6 in (*S*)-MeCPP-115–treated mice (p=0.0002) vs. vehicle treatment (**Fig. 4H**; treatment: F(1,90)=7.734, p=0.0066; time: F(8,90)=4.692, p<0.0001; interaction: F(8,90)=3.175, p=0.0032).

**Figure 4.**
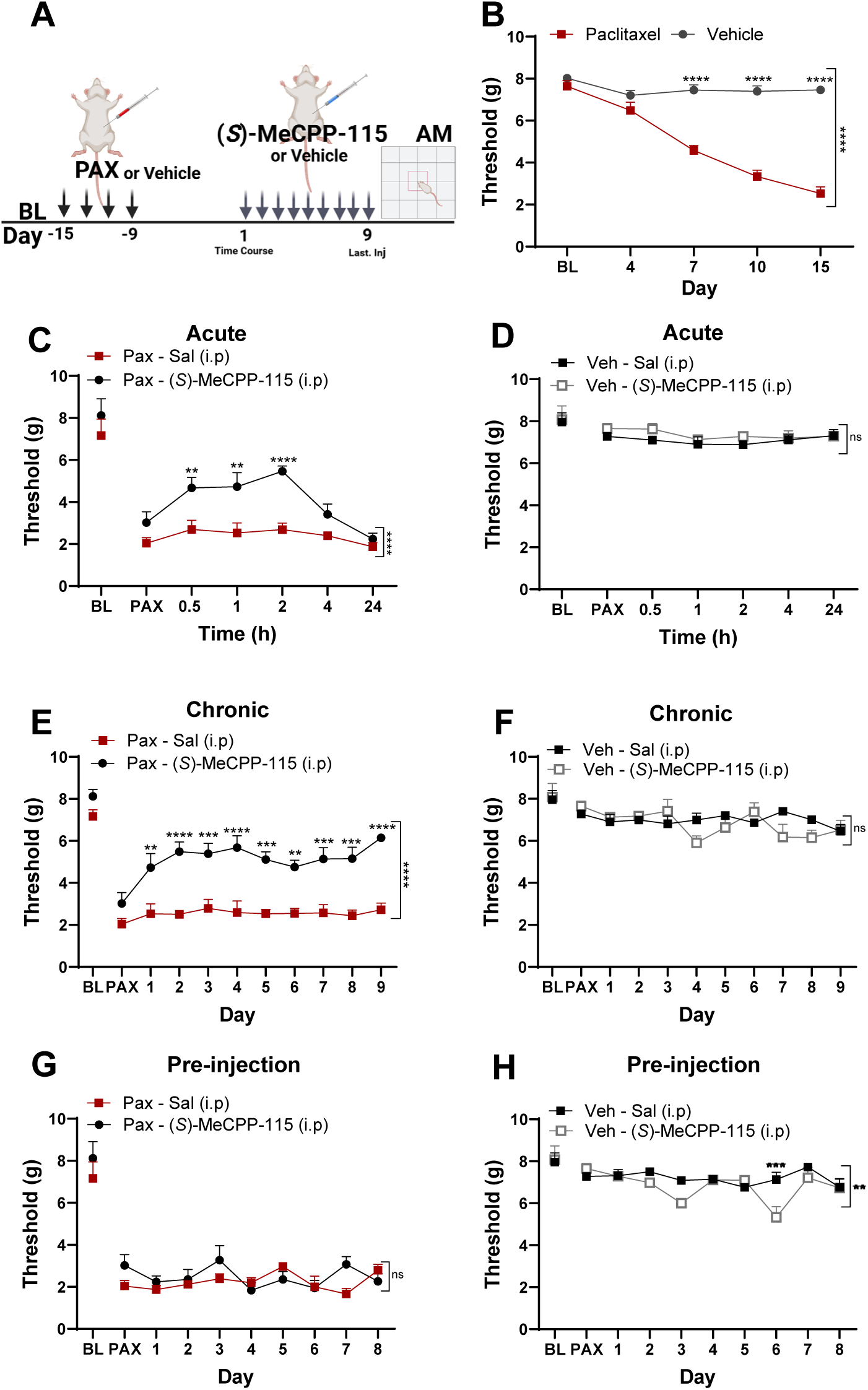
Systemic (*S*)-MeCPP-115 alleviates paclitaxel-induced mechanical hypersensitivity in mice. Experimental design (**A**). Paclitaxel reduced paw withdrawal thresholds compared with vehicle-treated animals (**B**). Acute systemic administration of (*S*)-MeCPP-115 (10 mg/kg, i.p.) in paclitaxel-treated mice induced a significant attenuation of aberrant mechanical hypersensitivity(**C**), whereas no change in paw withdrawal thresholds was observed in control mice (**D**). Repeated i.p. dosing with (*S*)-MeCPP-115 (10 mg/kg, i.p.) for 9 consecutive days progressively attenuated paclitaxel-induced mechanical hypersensitivity (**E**) without altering paw withdrawal thresholds in vehicle-treated animals (**F**). The antiallodynic action (*S*)-MeCPP-115 was not sustained 24 h after the final dose (**G**) and remained ineffective in controls receiving the same doses of (*S*)-MeCPP-115 (**H**). Data are expressed as mean ± SEM (n = 6 per group). *p* < 0.05, **p** < 0.01, ***p*** < 0.001 vs. respective controls (two-way ANOVA followed by Sidak’s post hoc test).

Locomotor activity was assessed immediately after von Frey testing in mice that received chronic i.p. treatments. (*S*)-MeCPP-115 did not reliably alter total distance traveled (**Fig. 5A**; F(3,20)=1.348, p=0.2871) or horizontal activity (**Fig. 5B**; F(3,20)=1.125, p=0.3626). Similarly, no changes were observed in rest time (**Fig. 5C**; F(3,20)=1.476, p=0.2513) or movement time (**Fig. 5D**; F(3,20)=1.476, p=0.2513). Anxiety-related behavior was not observed with (*S*)-MeCPP-115 treatment, as indicated by the absence of differences in time spent in the center of the arena (**Fig. 5E**; F(3,20)=1.510, p=0.2426). Finally, ambulatory velocity remained unchanged across groups (**Fig. 5F**; F(3,20)=1.023, p=0.4035), (**Fig. 5A–F**; all p>0.24).

**Figure 5.**
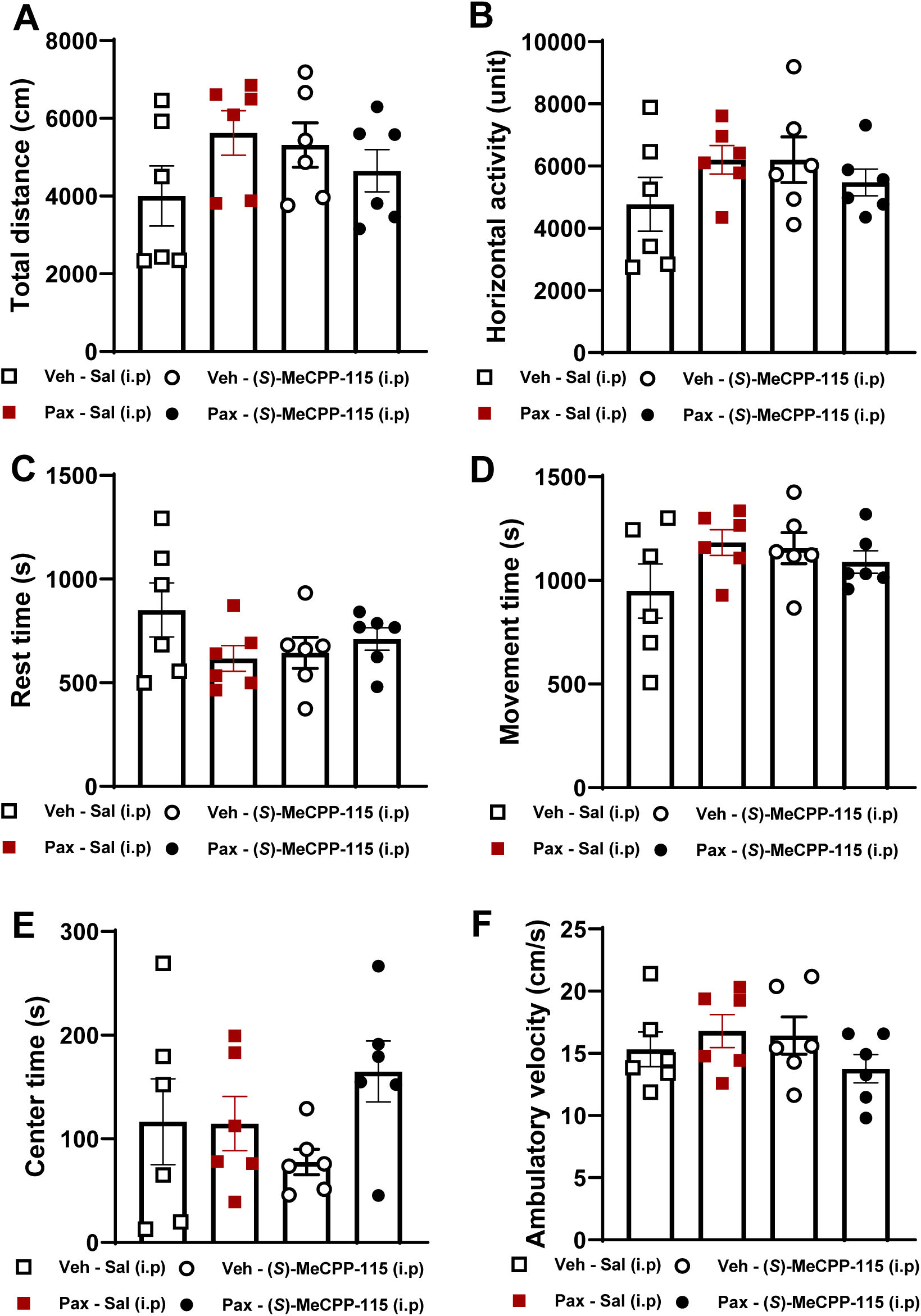
Systemic (*S*)-MeCPP-115 does not alter locomotor activity in paclitaxel-treated mice. Locomotor test conducted 1 h after the final day of dosing with (*S*)-MeCPP-115 (10 mg/kg/day x 9 days, i.p.) or saline (i.p.) administration. No significant differences were found among groups in total distance traveled (**A**), horizontal activity (**B**), rest time (**C**), movement time (**D**), time spent in the center (**E**), or ambulatory velocity (**F**). Data are expressed as mean ± SEM (n = 6 per group). Statistical analysis: one-way ANOVA followed by Tukey’s post hoc test.

### Intrathecal administration of (*S*)-MeCPP-115 reduces paclitaxel-evoked mechanical hypersensitivity with minor motor effects

Following induction of paclitaxel-induced mechanical hypersensitivity as performed in the previous cohort, mice received seven daily i.t. injections of (*S*)-MeCPP-115 (100 µg, i.t.) or vehicle **(Fig. 6A)**. Paclitaxel-treated mice developed mechanical hypersensitivity (**Fig. 6B**; treatment: F(4,110)=14.90, p<0.0001; time: F(4,110)=15.73, p<0.0001; interaction: F(1,110)=137.4, p<0.0001), which appeared by day 4 (p=0.0029) and persisted through day 15 (p<0.0001) following initiation of paclitaxel dosing. A single (*S*)-MeCPP-115 injection (100 µg, i.t.) reduced paclitaxel-induced mechanical hypersensitivity for over 4h, with effects no longer observed by 24 h post i.t. injection (**Fig. 6C**; interaction: F(5,60)=5.843, p=0.0002; time: F(5,60)=3.298, p=0.0107; treatment: F(1,60)=45.61, p<0.0001). Vehicle controls showed no change in mechanical paw withdrawal thresholds over time following acute i.t. injection (**Fig. 6D**; interaction: F(5,60)=1.948, p=0.0997; time: F(5,60)=3.029, p=0.0167; treatment: F(1,60)=0.1436, p=0.7061). Furthermore, (*S*)-MeCPP-115 consistently reversed paclitaxel-evoked hypersensitivity at 1h post-injection under repeated daily i.t. dosing (**Fig. 6E**; interaction: F(7,80)=3.878, p=0.0011; time: F(7,80)=1.576, p=0.1547; treatment: F(1,80)=133.8, p<0.0001), consistent with a spinal site of action. Paw withdrawal thresholds in saline controls remained stable following repeated i.t. dosing with (*S*)-MeCPP-115 (**Fig. 6F**; interaction: F(7,79)=0.9355, p=0.4841; time: F(7,79)=2.876, p=0.0100; treatment: F(1,79)=0.02340, p=0.8788). No residual antiallodynic effect was detected 24h after dosing (**Fig. 6G**; interaction: F(6,70)=0.5271, p=0.7859; time: F(6,70)=2.250, p=0.0483; treatment: F(1,70)=0.01006, p=0.9204), and thresholds in vehicle controls were also unchanged (**Fig. 6H**; interaction: F(6,70)=1.272, p=0.2814; time: F(6,70)=2.350, p=0.0399; treatment: F(1,70)=0.1894, p=0.6647).

**Figure 6.**
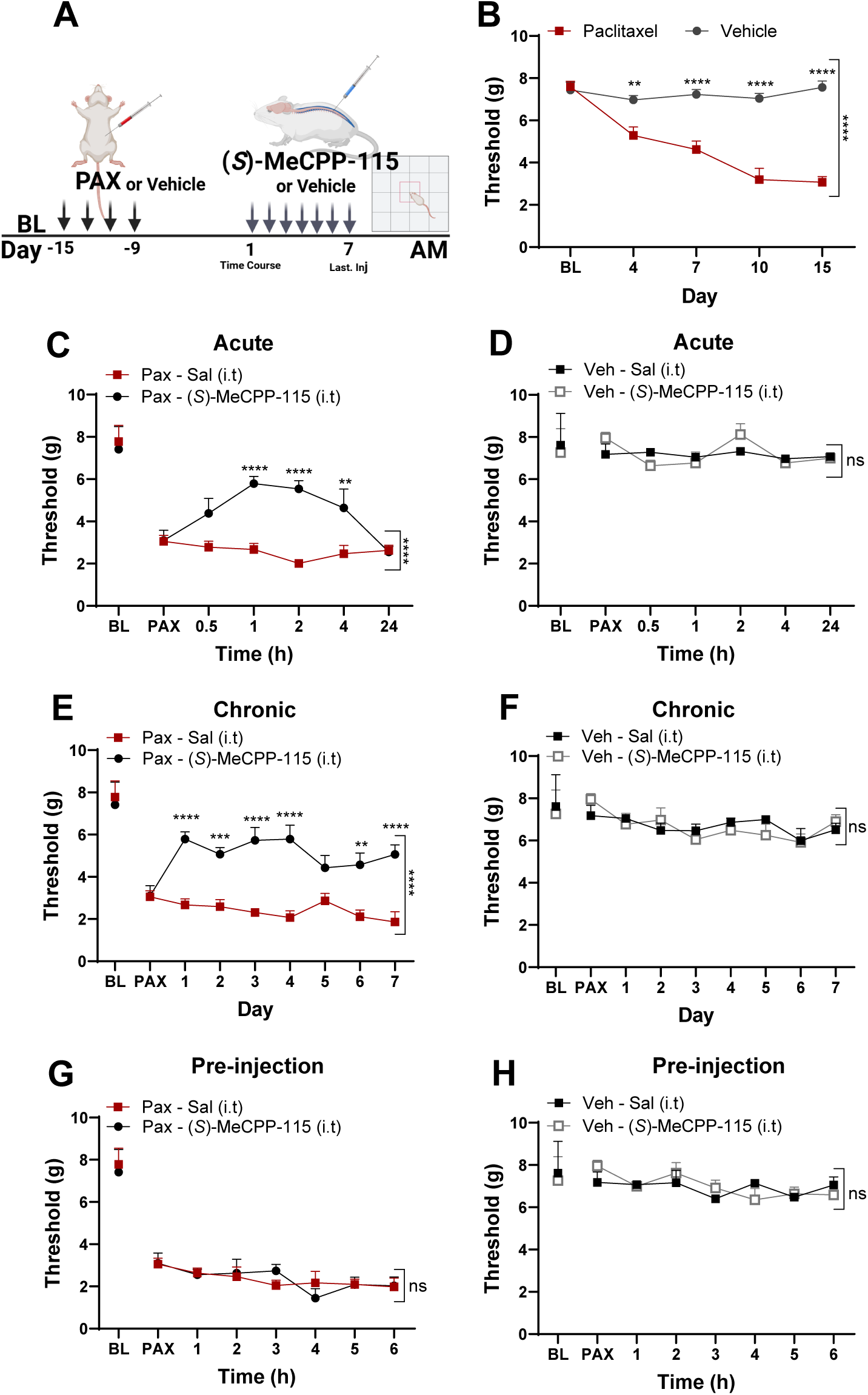
Intrathecal (*S*)-MeCPP-115 reverses paclitaxel-induced mechanical hypersensitivity in mice. Schematic representation of the experimental timeline (**A**). Paclitaxel markedly decreased paw withdrawal thresholds compared with vehicle-treated mice (**B**). Acute intrathecal delivery of (*S*)-MeCPP-115 (100 µg, i.t.) elicited antiallodynic effects lasting greater than 4 hours in paclitaxel-treated animals (**C**) but produced no changes in paw withdrawal thresholds in control mice that did not receive paclitaxel (**D**). Repeated intrathecal treatment with (*S*)-MeCPP-115 (100 µg, i.t.) for 7 consecutive days elevated mechanical paw withdrawal thresholds in paclitaxel-treated mice (**E**) without altering paw withdrawal thresholds in vehicle-treated mice that did not receive paclitaxel (**F**). The anti-allodynic effect of (*S*)-MeCPP-115 (100 µg/day x 7 days, i.t.) dissipated 24 h after the last administration (**G**) and remained absent in controls that receive vehicle in lieu of paclitaxel (**H**). Data are expressed as mean ± SEM (n = 6 per group). *p* < 0.05, **p** < 0.01, ***p*** < 0.001 vs. respective controls (two-way ANOVA followed by Sidak’s post hoc test).

Locomotor activity was assessed after von Frey testing in paclitaxel and Cremophor vehicle treated mice that received chronic i.t. dosing with (*S*)-MeCPP-115 or its vehicle (**Fig. 7A–F**). Intrathecal (*S*)-MeCPP-115 did not alter total distance traveled (**Fig. 7A**; F(3,19)=1.600, p=0.2224). Horizontal activity showed a treatment effect (**Fig. 7B**; F(3,19)=5.890, p=0.0051), with vehicle-treated mice receiving (*S*)-MeCPP-115 showing lower horizontal activity compared with both paclitaxel-treated mice that received (i.t.) saline (mean difference = 1853 units; 95% CI: 287.2–3419; p=0.0170) and vehicle treated mice that received (i.t.) saline (mean difference = 1926 units; 95% CI: 360.2–3492; p=0.0128). Vehicle-treated mice receiving (*S*)-MeCPP-115 (i.t.) also exhibited increases in rest time (**Fig. 7C**; F(3,19)=4.024, p=0.0225) and decreases in movement time (**Fig. 7D**; F(3,19)=4.024, p=0.0225) relative to paclitaxel-treated controls that received 7 consecutive once daily (i.t.) injections of saline. Post hoc comparisons failed to reveal any other reliable differences between groups. Time spent in the center (**Fig. 7E**; F(3,19)=0.7130, p=0.5562) and ambulatory velocity (**Fig. 7F**; F(3,19)=1.592, p=0.2244) were unaffected by any treatment.

**Figure 7.**
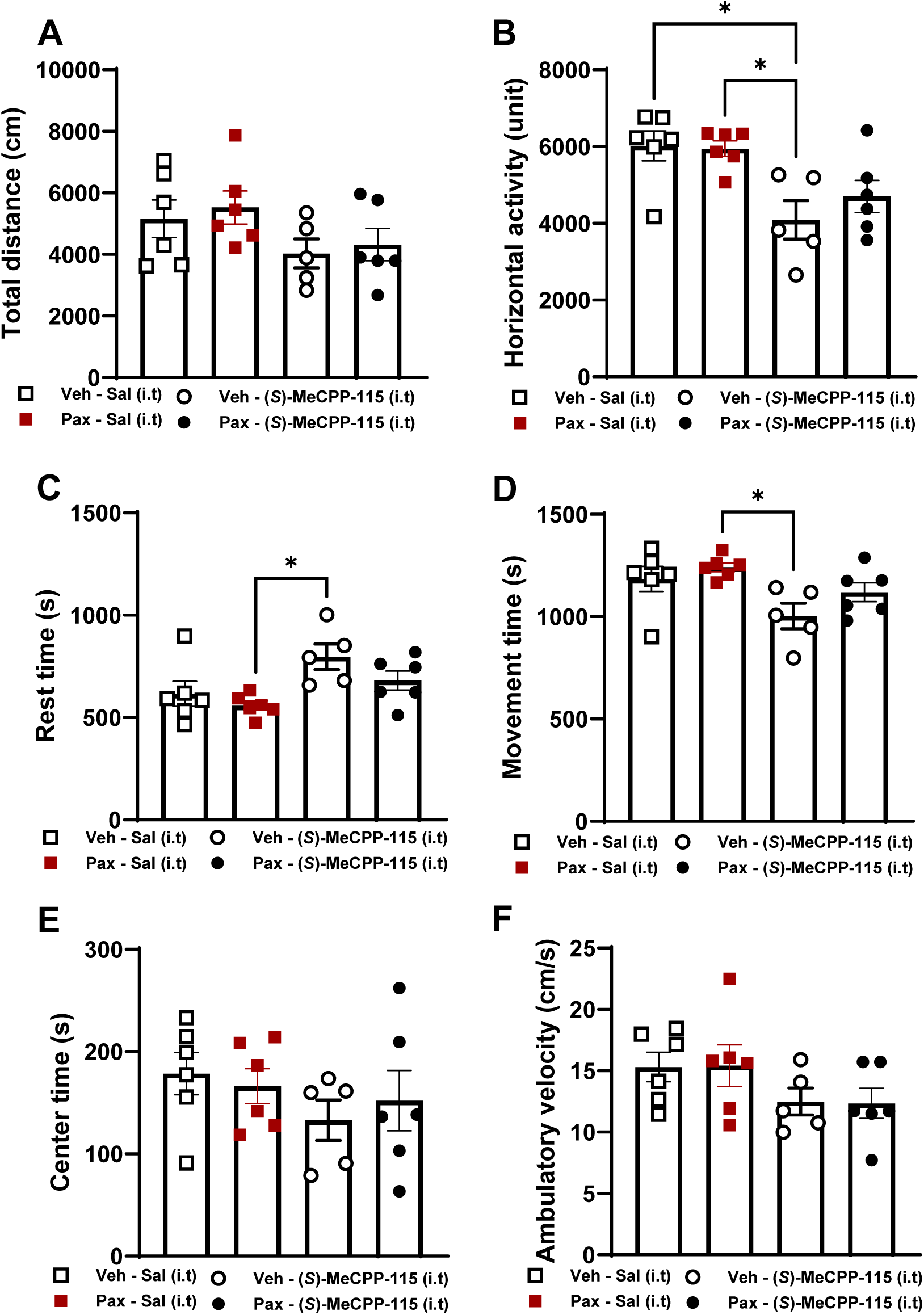
Effects of repeated intrathecal injection of (*S*)-MeCPP-115 and vehicle on locomotor activity. (*S*)-MeCPP-115 (100 µg, i.t.) administered once daily over 7 consecutive days did not alter total distance traveled in paclitaxel-or vehicle-treated mice (**A**). Minor but statistically significant alterations were observed in horizontal activity (**B**), rest time (**C**), and movement time (**D**), suggesting mild modulation of locomotor activity in vehicle-treated mice that received (*S*)-MeCPP-115 (100 µg, i.t. x 7 days). No differences were observed between groups in time spent in the center of the activity meter (**E**) or in ambulatory velocity (**F**). Data are expressed as mean ± SEM (n = 6 per group). Statistical analysis: one-way ANOVA followed by Tukey’s post hoc test.

## Discussion

The present study identifies (*S*)-MeCPP-115, a selective GABA-AT inactivator developed from CPP-115, as an effective analgesic candidate in a mouse model of paclitaxel-induced peripheral neuropathy. Before assessing its antinociceptive effects *in vivo*, we first confirmed *in vitro* that (*S*)-MeCPP-115 does not interfere with paclitaxel’s cytotoxicity in a 4T1 breast cancer cell line. This is a key requirement for any analgesic candidate for CIPN, as preserving chemotherapy efficacy is paramount. (*S*)-MeCPP-115 reduced established mechanical hypersensitivity following both i.p. and i.t. routes of administration. Furthermore, efficacy was maintained following repeated dosing with no signs of tolerance. Moreover, (*S*)-MeCPP-115 did not alter mechanical paw withdrawal thresholds in control mice that did not receive paclitaxel following acute or chronic dosing via either systemic (i.p.) or intrathecal (i.t.) routes of administration. The rationale for GABA-AT inhibition stems from evidence that CIPN involves a loss of inhibitory tone within spinal and supraspinal nociceptive circuits. Our studies confirm that (*S*)-MeCPP-115 acts, at least in part, through a spinal site of action. Paclitaxel-induced reductions in GABAergic inhibition have been observed in the anterior cingulate cortex, and thalamus as well as the spinal cord. Approaches that restore GABA function such as GABA transporter blockade or GABAergic precursor transplantation in the spinal cord attenuate neuropathic pain in preclinical models [5, 14–16]. By preventing GABA degradation, GABA-AT inactivators offer an indirect but sustained means of increasing inhibitory signaling without the adverse effects associated with direct GABA receptor agonists or benzodiazepines [17–19].

Our group previously reported that OV329, a highly potent GABA-AT inactivator, suppressed pain behaviors in models of neuropathic and inflammatory pain in male and female mice without inducing conditioned place preference or self-administration [6]. (*S*)-MeCPP-115, although less potent than OV329, displays markedly greater selectivity for GABA-AT over ornithine aminotransferase (OAT), which may confer improved tolerability. In a CIPN model, (*S*)-MeCPP-115 produced robust suppression of paclitaxel-induced mechanical hypersensitivity without inducing tolerance. Both systemic and intrathecal administration reversed paclitaxel-induced hypersensitivity, although effects dissipated within 24 hours. Importantly, locomotor impairment was minimal after repeated intrathecal dosing and was absent following 9 days of repeated intraperitoneal treatment. From a translational perspective, patients requiring intrathecal therapy typically receive continuous infusion via implanted pumps. Therefore, repeated daily intrathecal injections in mice, which are more stressful than intraperitoneal administration, do not accurately reflect the clinical scenario [20]; the absence of an implanted pump system thus represents a limitation of this study.

In conclusion, (*S*)-MeCPP-115 did not interfere with antitumor activity *in vitro* and effectively reduced paclitaxel-induced mechanical hypersensitivity without producing tolerance or major motor side effects *in vivo*. These results reinforce the potential of enhancing GABAergic signaling through GABA-AT inhibition as a promising strategy for the treatment of CIPN. This mechanism may offer advantages in terms of selectivity and safety compared to conventional approaches to GABA modulation. While (*S*)-MeCPP-115 stands out as an initial example of a next-generation inhibitor, future studies should prioritize broader pharmacological characterization, evaluation of sex differences, exploration of additional pain models, and investigation of long-term safety to consolidate its translational potential.

## Acknowledgments

This work was supported by the National Institutes of Health [Grants DA030604 (to R.B.S.), NS123057 (to R.B.S. with subcontract to A.G.H.), and DA047858 (to A.G.H.)]

## References

1. Lee, K.T., et al., Chemotherapy-Induced Peripheral Neuropathy (CIPN): A Narrative Review and Proposed Theoretical Model. Cancers (Basel), 2024. 16(14).

2. Peek, A.L., et al., Brain GABA and glutamate levels across pain conditions: A systematic literature review and meta-analysis of 1H-MRS studies using the MRS-Q quality assessment tool. Neuroimage, 2020. 210: p. 116532.

3. Ferrier, J., et al., Cholinergic Neurotransmission in the Posterior Insular Cortex Is Altered in Preclinical Models of Neuropathic Pain: Key Role of Muscarinic M2 Receptors in Donepezil-Induced Antinociception. J Neurosci, 2015. 35(50): p. 16418–30.

4. Nashawi, H., et al., Paclitaxel Causes Electrophysiological Changes in the Anterior Cingulate Cortex via Modulation of the gamma-Aminobutyric Acid-ergic System. Med Princ Pract, 2016. 25(5): p. 423–8.

5. Qian, X., et al., Current status of GABA receptor subtypes in analgesia. Biomed Pharmacother, 2023. 168: p. 115800.

6. Wirt, J.L., et al., Efficacy of GABA aminotransferase inactivator OV329 in models of neuropathic and inflammatory pain without tolerance or addiction. Proc Natl Acad Sci U S A, 2025. 122(1): p. e2318833121.

7. Chan, K., et al., Vigabatrin-Induced Retinal Functional Alterations and Second-Order Neuron Plasticity in C57BL/6J Mice. Invest Ophthalmol Vis Sci, 2020. 61(2): p. 17.

8. Hawker, M.J. and N.J. Astbury, The ocular side effects of vigabatrin (Sabril): information and guidance for screening. Eye (Lond), 2008. 22(9): p. 1097–8.

9. Juncosa, J.I., et al., Design and Mechanism of (S)-3-Amino-4-(difluoromethylenyl)cyclopent-1-ene-1-carboxylic Acid, a Highly Potent gamma-Aminobutyric Acid Aminotransferase Inactivator for the Treatment of Addiction. J Am Chem Soc, 2018. 140(6): p. 2151–2164.

10. Pan, Y., et al., *(1S*, *3S)-3-amino-4-difluoromethylenyl-1-cyclopentanoic acid (CPP-115), a potent gamma-aminobutyric acid aminotransferase inactivator for the treatment of cocaine addiction*. J Med Chem, 2012. 55(1): p. 357–66.

11. Doumlele, K., et al., *A case report on the efficacy of vigabatrin analogue (1S*, *3S)-3-amino-4-difluoromethylenyl-1-cyclopentanoic acid (CPP-115) in a patient with infantile spasms*. Epilepsy Behav Case Rep, 2016. 6: p. 67–9.

12. Feja, M., et al., OV329, a novel highly potent gamma-aminobutyric acid aminotransferase inactivator, induces pronounced anticonvulsant effects in the pentylenetetrazole seizure threshold test and in amygdala-kindled rats. Epilepsia, 2021. 62(12): p. 3091–3104.

13. Moschitto, M.J. and R.B. Silverman, Synthesis of (S)-3-Amino-4-(difluoromethylenyl)-cyclopent-1-ene-1-carboxylic Acid (OV329), a Potent Inactivator of gamma-Aminobutyric Acid Aminotransferase. Org Lett, 2018. 20(15): p. 4589–4592.

14. Vaysse, L., et al., GABAergic pathway in a rat model of chronic neuropathic pain: modulation after intrathecal transplantation of a human neuronal cell line. Neurosci Res, 2011. 69(2): p. 111–20.

15. Jergova, S., et al., Intraspinal transplantation of GABAergic neural progenitors attenuates neuropathic pain in rats: a pharmacologic and neurophysiological evaluation. Exp Neurol, 2012. 234(1): p. 39–49.

16. Senba, E. and K. Kami, Potentiation of spinal GABA inhibition as a therapeutic target for chronic neuropathic pain: from transplantation to physical exercise. Ann Palliat Med, 2020. 9(5): p. 2430–2436.

17. Vgontzas, A.N., A. Kales, and E.O. Bixler, Benzodiazepine side effects: role of pharmacokinetics and pharmacodynamics. Pharmacology, 1995. 51(4): p. 205–23.

18. Thompson, S.M., Modulators of GABA(A) receptor-mediated inhibition in the treatment of neuropsychiatric disorders: past, present, and future. Neuropsychopharmacology, 2024. 49(1): p. 83–95.

19. Goldschen-Ohm, M.P., Benzodiazepine Modulation of GABA(A) Receptors: A Mechanistic Perspective. Biomolecules, 2022. 12(12).

20. Arman, A. and M.R. Hutchinson, Intrathecal implantation surgical considerations in rodents; a review. J Neurosci Methods, 2021. 363: p. 109327.

